# Cortical Field Model of Complex Spiral Traveling Waves

**DOI:** 10.64898/2026.01.06.698037

**Authors:** Ghanendra Singh

## Abstract

Complex spiral traveling waves observed experimentally occur across the cortex. The underlying mechanisms responsible for generating such mesoscopic activity are not well understood. Understanding how local cortical neuronal populations interact to produce emergent spiral dynamics during cognitive processing remains unknown. Therefore, to bridge this gap, a spatiotemporal cortical field rate model of local cortical circuits, composed of excitatory and three distinct time-scale inhibitory populations, is proposed. This model is extended to a two-dimensional cortical sheet, consisting of both nonlinear local interactions and diffusive global coupling, with distance-dependent axonal delays. Simulation results indicate mixed-mode oscillations occur in the local circuits, which may represent the coexistence of multiple rhythms and show the emergence of complex dynamics, such as rotating spirals with annihilation events. Spiral waves differentially respond to the strength of the grating input stimulus and exhibit working memory-like characteristics. Also hypothesize that local patterns, such as planar, source, sink, or concentric across the cortex, might be an inherent integral part of the spiral state dynamics.

## Introduction

Spiral waves exist in the nature and aggregate slime *Dictyostelium* mounds by complex propagating modes across prestalk and prespore cells with different excitability or oscillatory properties (Early et al., 1995; Siegert & Weijer, 1995). Scroll waves in three dimension at prestalk and planar waves at prespore regions organise this slug(Siegert & Weijer, 1992). Interestingly, behavioural movements also modifies the geometry of these patterns. Earlier models clearly showed that cAMP waves propagation undergoes transition from spiral to scroll waves in 3D formation of mound (Vasiev & Weijer, 1999). Numerous evidence of spiral waves such as in the heart (Holden, 1997),in turtle visual cortex (Prechtl et al., 1997), in mammalian rat cortex with dominant excitatory interactions (Huang et al., 2004), traveling waves in the monkey motor cortex (Rubino et al., 2006), across the mouse neocortex during sleep-like states (Huang et al., 2010), suppressive traveling waves in awake monkey (Chemla et al., 2019), in visual cortex (Sato et al., 2012), in human cortex (Muller et al., 2018), and in mouse cortex (Liang et al., 2023). Sleep spindles also seen as rotating (Muller et al., 2016) or spiral waves for memory consolidation (Y. Xu et al., 2025). Further, many different patterns exists as sources, sinks, and saddles during local traveling wave interactions (Liang et al., 2021). Locally, during the wakeful state transition, the number of sources, sinks, and saddles increased, representing multiple brain states (Liang et al., 2023). In essence, these co-existing complex patterns might act as distributed parallel *spiral computing units* in a dynamically changing state space (Gong & Van Leeuwen, 2009) across the cortical grid layered architecture.

Spiral traveling waves are distributed across the cortex as an organizing principle for neuronal activity (Ye et al., 2023). These spatiotemporal patterns (Schiff et al., 2007) evolves during working memory (Bhattacharya et al., 2022). Traveling waves are spontaneous in nature and undergoes collision while propagating toward each other also results in annihilation events (Wu et al., 1999). Direction of traveling waves also regulates memory processing (Mohan et al., 2024). With three different inhibitory populations, there is a substantial inhibitory role, as also observed in LFP recordings (Teleńczuk et al., 2017). Feedback and feedforward connections shape the traveling waves in feature-selective motifs. Generally, excitatory coupling dominates in local interactions but weak inhibitory coupling might enable long-distance spiral wave generation across the cortex. Such delayed excitation from distal weakly coupled local cortical regions might result in the propagation of traveling waves in the cortex (G. B. Ermentrout & Kleinfeld, 2001). Earlier, the spatial version of the Wilson-Cowan model (Wilson & Cowan, 1972, 1973) theorized about traveling waves. Therefore, its worth investigating the local and global inhibitory coupling occuring at different time scales due to multiple inhibitory population motifs (Campagnola et al., 2022).

### Inhibition role in the cortical oscillations

Interneurons are classified into different subtypes based on the expression of parvalbumin (PV), somatostatin (SOM), or vasointestinal peptide (VIP), which divides them into three groups (Rudy et al., 2011), which mainly cover all interneurons (Tremblay et al., 2016). PV and SOM neurons most strongly inhibit pyramidal cells. SOM neurons mostly integrate excitation from the local population (Adesnik et al., 2012), whereas PV neurons inhibit the cell body, while SOM neurons inhibit the distal dendritic arbor, shaping differential integration by pyramidal neurons (Tremblay et al., 2016). SOM neurons inhibit PV neurons (Pfeffer et al., 2013), regulating the inhibition mode of PV neurons. Different inhibitions modulate the cortical wave dynamics (Xiao et al., 2012). Early and late traveling waves exist across the barrel cortex of the mice with four distinct patterns (irregular, plane, ring and spiral) that emerge in tangential slices and their interactions might underlie the mechanisms of computation or information processing (Huang et al., 2004). Earlier spiking (Keane & Gong, 2015) models were used to describe the dynamic propagation of these traveling waves but how does the distinct inhibitory neuronal populations plays a role in the occurrence of these time dependent complex wave patterns is unclear.

Long-range connections efficiently transfer information between cortical regions in the wakeful state and interactions between multiple, coexisting localized wave patterns enable spiral wave annihilation-based dynamical computations to be implemented in a distributed and parallel manner (Zhang et al., 2014). Macroscopic brain states have an affect on the local wave patterns (Liang et al., 2023). What dynamical mechanisms relates the mesocopic activity with macroscopic states? Earlier theoretical prediction of distributed dynamic computations (Gong & Van Leeuwen, 2009) points to these complex spiral traveling waves based computations. A mean field model described the role of inhibition in cortical wave networks (Palkar et al., 2023). Next, inhibition could enable information to be readily transmitted across distances without the neural activity blowing up. PV, SOM, and VIP neurons contributes to disynaptic inhibition and topdown modulation (Zhang et al., 2014). These cortical circuits, with multiple inhibitory neurons, could also perform predictive coding via oscillations (Lee et al., 2025). Recently proposed spiking network model (Sarkar & Ermentrout, 2025) described co-existence of two different rhythms with large and small amplitudes occurring in the same cortical circuit composed of two different interneurons, PV and SOM. Interestingly, it refers to a similar phenomenon of mixed mode oscillations (MMOs) (Desroches et al., 2012) well described with large amplitude oscillations (LAOs) and small amplitude oscillations (SAOs) existing in the slow-fast dynamical regime. In summary, MMOs naturally fits as a robust mathematical framework to theoretically study the properties of complex spiral traveling waves occuring in the brain cortex.

### Mean field population model of the cortex

Coupled oscillator model can be utilized to generate complex spatiotemporal patterns, including planar, spiral, and irregular waves in the cortex. Simplified rate models can also be used to understand the significance of distinct inhibitory neuronal population activity regimes of VIP, PV and SOM. Then why specifically a mean field model is utilized in this study instead of spiking network model? There exist a strong evidence for choosing the mean firing rate model over the individual spiking network model is that these neocortical distinct inhibitory populations, such as somatostatin (SOM) or vasoactive intestinal peptide (VIP), are active as populations rather than individual neurons and work as cooperative units to provide a strong nonlinear response in the cortical circuit. (Karnani et al., 2016). I took inspiration from this proposed cooperative model based on the experimental findings to understand the mesoscopic dynamics. Population-based local excitation or disinhibition might underlie inhibition co-activity. SOM or VIP interneurons fire together and recruit additional cells, which further amplify the population’s output. Hence, these different inhibitory populations could function as cooperative processing units to generate an amplifying nonlinearity in their circuit output.

Distinct interneurons differentially regulate a cortical local circuit which could be involved in the cognitive processes such as encoding, maintenance, or retrieval of stimuli during working memory (Wang et al., 2004). Also, proposed a framework to demonstrate that interneuron subtypes exert differential inhibition. Distinct role of PV and SST interneurons exist in the working memory (Kim et al., 2016). SOM activity is increased during the delay period, suggesting its role in the maintenance phase of working memory. They show that ensembles of specific types of inhibitory interneurons generate coordinated activity in the mouse visual cortex. Diverse interneuron ensembles to function as distinct operational units can exhibit synchronous activity. Overall, the above studies supports the arguement that population activity based model could explain certain aspects of the experimentally observed rotating complex spiral traveling waves.

### Heterogeneity of inhibitory timescales

Multiple timescales exists at the level of single neurons or neuronal populations within a cortical region. Stable population-level working memory representations exist in pre-frontal cortex (PFC) despite strong temporal neural dynamics (Murray et al., 2017). Distinct inhibitory coupling or disinhibitory mechanisms that exist in the local cortical circuits across different layers might have a role in working memory. Cortical neurons exhibit multiple intrinsic timescales related to the spontaneous dynamics. In (Trepka et al., 2024), provide evidence for plastic, finegrained gradients of timescales within PFC that can influence both single-cell and population coding, pointing to the importance of these different timescales.

What could be the underlying mechanisms for multiple time scales (Spitmaan et al., 2020)? Inhibitory neuron subtypes perform context-dependent modulation of excitatory activity, as well as regulate the experience-dependent plasticity of excitatory circuits (Hattori et al., 2017). Local spiking patterns shape cortical traveling waves (Ye et al., 2023). Described a unifying routing framework for macroscopic brain activity via traveling waves reconfiguration through direction changes, where spiral, sources, and sinks are local patterns as state variables for the cortex (Vinao-Carl et al., 2025). Does multiple time scales play a role in building an association between short-term and long-term memories (Szatmáry & Izhikevich, 2010)? How does a population effectively utilize a whole distribution of timescales provided by distinct inhibitory neurons for the encoding, maintenance, and retrieval of items in the working memory (Cavanagh et al., 2020)? Hence, I argue that these distinct timescales builds local circuits with slow-fast dynamical regimes that effectively multiplexes different rhythms using mixed-mode oscillations framework.

### Simple vs complex spiral traveling waves

Simple traveling waves occurs in the cortex typically exhibit smooth, coherent propagation with a single dominant direction and stable velocity, reflecting low-dimensional, spatially organized population dynamics often observed during sleep, anesthesia, or sensory-evoked states (Muller et al., 2018; Townsend & Gong, 2018). In contrast, complex traveling waves show multidirectional flow, rapid changes in speed or trajectory, fragmentation of wavefronts, and interactions between multiple propagating motifs, producing richer spatiotemporal patterns linked to awake, active, and heterogeneous cortical states (Muller et al., 2016). These complex waves arise from nonlinear recurrent circuitry and structural heterogeneity at the mesoscale, allowing the cortex to flexibly route information and support dynamic coordination across distributed networks. I conducted numerical simulations of the four population neural field model to investigate the emergence of spatiotemporal dynamics under different input regimes and parameter configurations. The simulation results demonstrate that diverse activity patterns could arise from interactions between excitatory and inhibitory populations.

## Results

A spatially extended cortical mean field model is proposed, inspired by local cortical circuit connectivity in mice (Campagnola et al., 2022), incorporating one excitatory (PYR) and three inhibitory (PV, SOM, VIP) populations. Each population follows wilson–cowan rate dynamics (Wilson & Cowan, 1972, 1973). The model includes recurrent, feedback or feedforward loops to represent diverse intra and inter laminar circuit motif topologies (Campagnola et al., 2022). Earlier large-scale models of turtle visual cortex (Nenadic et al., 2003) with pyramidal and multiple inhibitory cells (Senseman & Robbins, 1999) resulted in propagating cortical waves with complex spatiotemporal structure. Further, rate based models have been effective in explaining neocortical responses to brief stimuli (Cowan et al., 2016).

Therefore, this study looked at five different aspects. First, local cortical circuit dynamics with two or three inhibitory populations producing oscillations along with bifurcation diagrams. Second, simulated the extended spatiotemporal cortical grid model that generates rich spatiotemporal activity including spiral waves, planar fronts, sources, sinks, and concentric patterns (Das et al., 2024). Third, annihilation events across the rotating spirals. Fourth, effect of the weak and strong grating stimulus on the 2d grid cortical field. Fifth, calculating phase gradient and phase velocity fields across excitatory and inhibitory cortical field.

### Local Cortical Circuits produces Normal and Mixed Mode Oscillations

Mixed-mode oscillations (MMOs) occur across chemical, biological, and neuronal systems, spanning multiple time scales and comprising fast and slow subsystems. A detailed review distinguishing the simple and complex MMOs and underlying mechanisms (canards, Shilnikov, or period-doubling) (Desroches et al., 2012).

This mathematical framework fits well to describe the underlying slow-fast mechanisms responsible for the coexistence of multiple rhythms across the cortex (Sarkar & Ermentrout, 2025). One can further investigate to establish the theory under-lying the relationship between local MMOs and complex spiral patterns.

Simulations in the figure 2 refer to the cortical circuit in fig. 2A. composed of excitatory PYR and inhibitory PV, and VIP. Normal oscillations with leading PV phase shown in fig 2B. and MMOs in fig. 2C. Bifurcation diagram is plotted in fig. 2D. Next, cortical circuit with VIP population shown in the figure 2E., with normal oscillations and MMOs in fig. 2F. and 2G along with bifurcation diagram in 2H. Bifurcation plots shows Hopf bifurcation (HB) and period doubling (PD) with small and large ranges.

**Figure 1:**
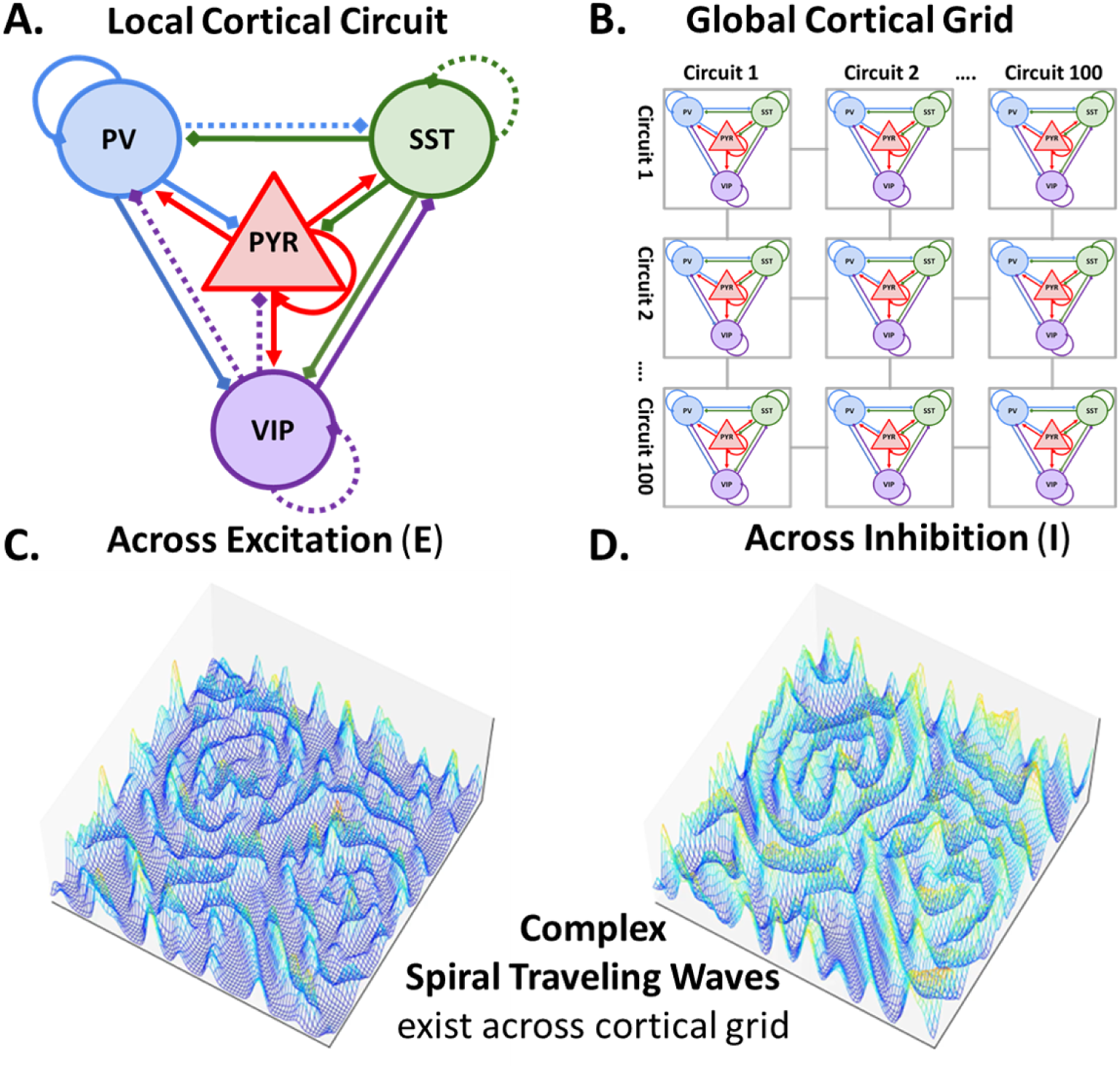
Complex spiral traveling waves across the cortical sheet. (A.) Local cortical circuit consisting of an excitatory pyramidal (PYR) population and three different types of inhibitory (SOM, PV, and VIP) population. Local connectivity architecture is based on (Campagnola et al., 2022) with strong interactions as solid lines for and weak interactions with dashed lines (B.) Global cortical 2D grid (100x100) composed of local cortical circuits coupled by diffusion representing the internal axonal delays between neuronal population. (C.) Complex spiral traveling waves field representing mean firing rate emerges across the 2D cortical grid of excitatory (E) and (D.) inhibitory population. These complex spiral traveling waves can be visualized in the supplementary video 1 and supplementary video 2 files.

**Figure 2:**
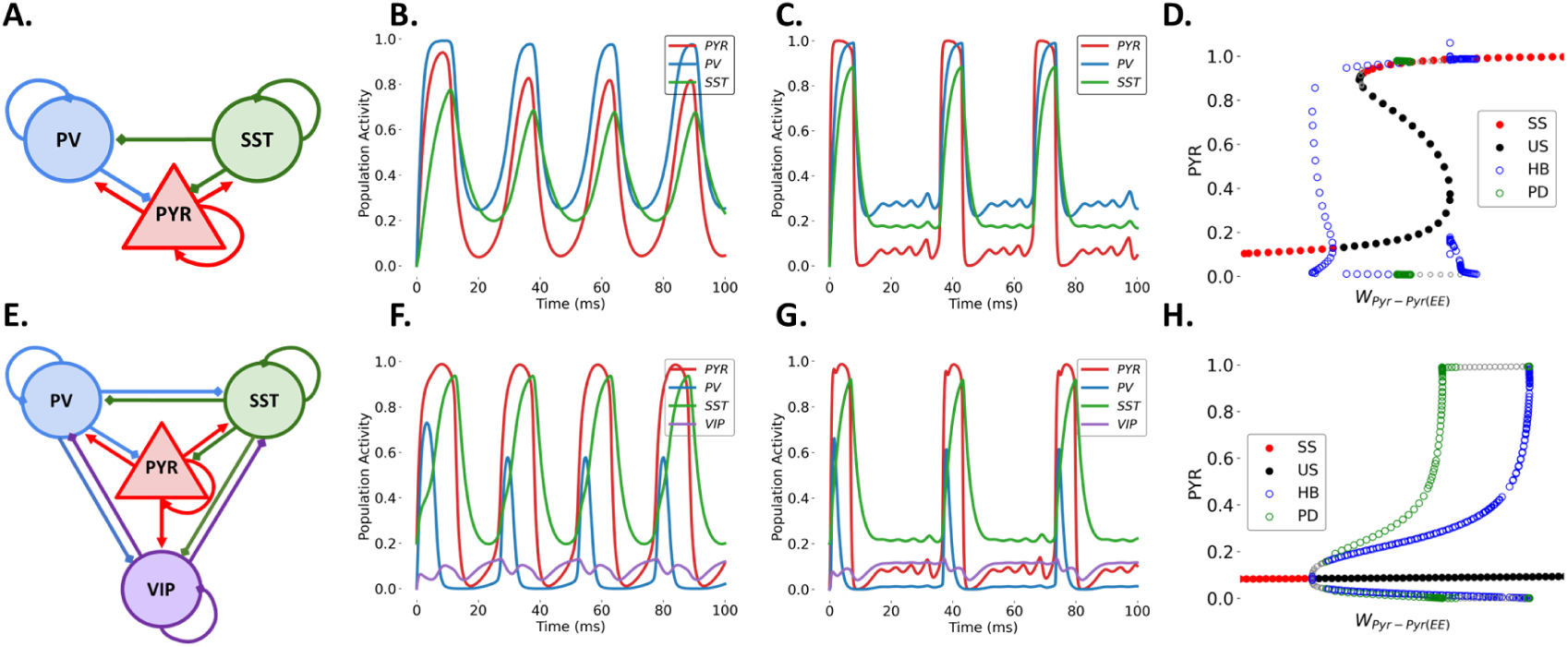
Local Cortical Circuit Dynamics with Normal and Mixed Mode Oscillations. (A.) Local cortical circuit with excitatory PYR and two inhibitory (PV, SST) populations. (B.) Normal oscillations of fast firing PV with leading phase, intermediate PYR and slow SST. (C.) Mixed mode oscillations (MMOs) occurs with fast PYR and slow inhibitory (PV and SST) population rates with periodic large amplitude oscillation (LAOs) of low frequency and small amplitude oscillations (SAOs) of high frequency. (D.) Bifurcation plots shows Hopf bifurcation (HB) and period doubling (PD) with narrow range for cortical circuit in A. (E.) Cortical circuit with addition VIP population. (F.) Normal oscillations and (G.) MMOs occurs across E. (H.) Bifurcation plots shows Hopf bifurcation (HB) and period doubling (PD) with wider range for cortical circuit in E. with three inhibitory population.

Different modes of MMOs through the phenomenon of PD that might be regulated by slow-changing synaptic variables or externally applied sensory inputs to local cortical circuits might explain the temporal interactions or co-existence of cortical rhythms during working memory (Miller et al., 2018). MMOs may provide narrow windows of robust interaction between slow and fast oscillations via unidirectional or bidirectional cross-frequency coupling (Hyafil et al., 2015) to store multiple items in working memory. So far, my objective was to focus on establishing a relationship between local cortical circuits and certain aspects of spiral patterns to appreciate the beauty of simple rate models in capturing the emergent complex traveling wave dynamics.

### Emergence of Complex Spiral Traveling Waves Patterns

Now, on extending the local cortical circuit into a 2d grid with inhibitory diffusive coupling to understand the mechanisms producing these complex patterns that play a critical role during cognitive tasks such as a working memory (Bhattacharya et al., 2022). Earlier proposed three layered 2D phenomenological computational model architecture of size 60x60 grid showed different spatiotemporal patterns such as planar, rotating waves, or radial (Bhattacharya et al., 2021). Simulation in the figure 3 produced these patterns with an example of a spiral core as tiny orange circle with low amplitude and two spiral arms in blue with high amplitude clearly visible in the green box (T2-T4), which confirms complex rotating spiral patterns as previously shown (Huang et al., 2004).

**Figure 3:**
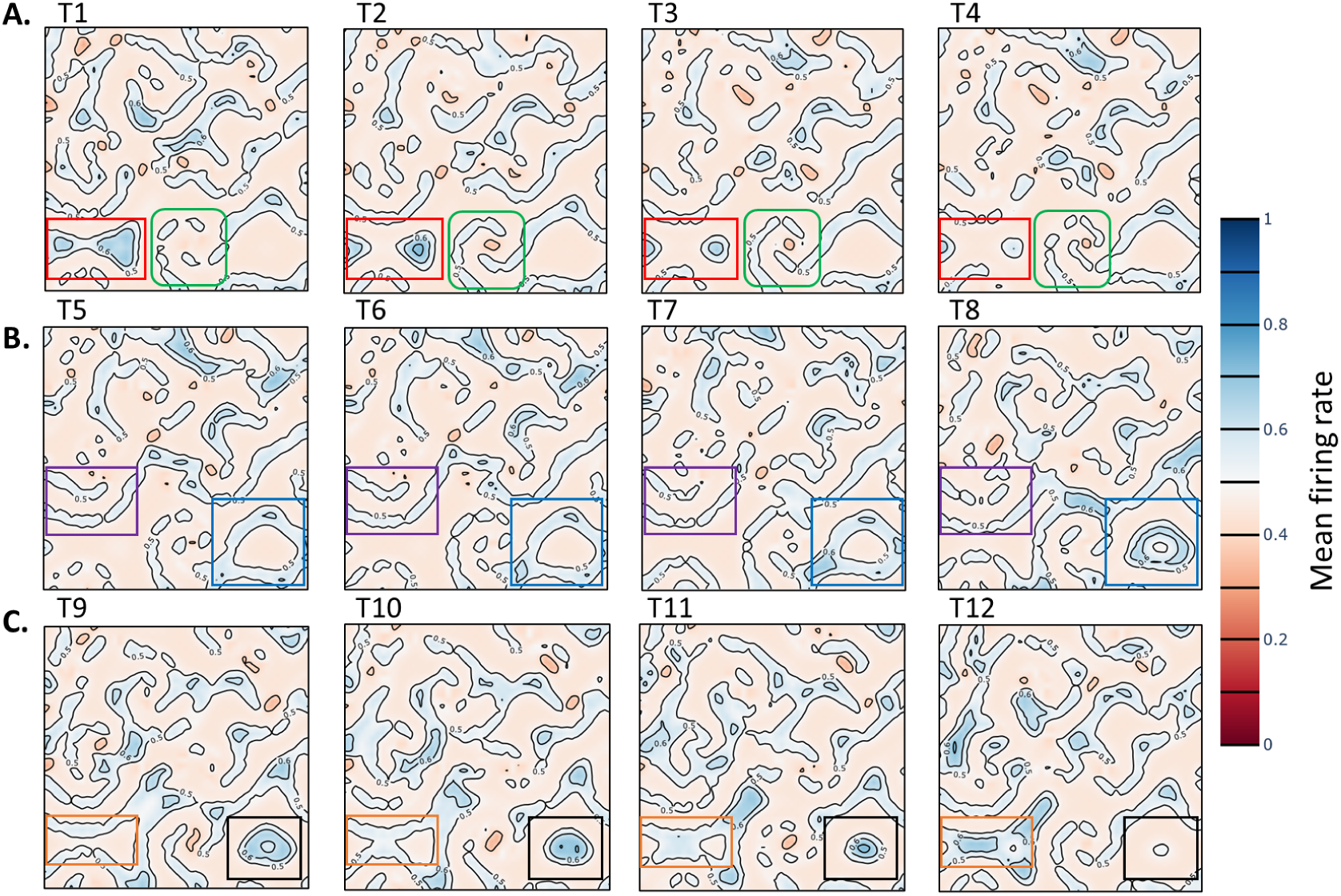
Propagation of different Complex Spiral Traveling Waves Patterns. Cortical field model displays various patterns simultaneous coexist together. (A.) Complex spiral pattern rotating clockwise shown in the green box varying with activity patterns. Adjacent to it is formation of sources converging to sink like patterns shown in (red). (B.) Activity patterns part of a spiral appear as planar propagating in a direction shown in the (purple) box. Adjacent activity shows converging concentric patterns as shown in the (blue) box. (C.) Propagating spiral patterns appearing as planer undergoes annihilation as shown in the (orange) box. Adjancent to it is sink like of patterns in (black).

Under moderate external input to the PYR, the model spontaneously develops spatially periodic activity patterns. In the figure 3, during the time steps (T1-T4), two local nodes appearing as sources and converging to sink like patterns (red box). A spiral pattern rotating clockwise, with an orange core at the center with low firing rate, and two arms with discrete activity patterns (green box). Next, during the time steps (T5-T8), planar patterns that might be part of the spirals appear to propagate in a particular direction (purple box). Concentric patterns appear to converge on a stable focus point (blue box). Furthermore, during the time steps (T9-T12), spiral arms, as planar patterns propagating towards each other, undergo annihilation (orange box). This annihilation event results in a slightly higher firing rate for a shorter duration. Lastly, sink-like patterns also appear (black box). In summary, simultaneous existence of different patterns across the cortical field observed both experimentally and from model predictions suggest that, in principle brain might be using such global complex spiral wave based computational mechanisms for efficient parallel information processing.

Spirals create more semi planar waves that move away from the stimulation site (Santos et al., 2014). I observed something similar that after the annihilation event of the spirals, semiplanar patterns move away from each other as seen in the figure 4. This validates the model predictions. Earlier it was asked that what could be the biological substrate for the generation of traveling waves (G. B. Ermentrout & Kleinfeld, 2001)? Now, I think, we might have some observations at the mechanistic and mesoscopic levels. Delayed excitations across different regions could be due to heterogeneity of inhibitory time scales across different layers. Stimulus with different frequencies resulted in harmonic responses which may suggest the observed period doubling phenomena in the model circuits. Connectivity of the population circuits in traveling waves might be distinct (Benucci et al., 2007).

**Figure 4:**
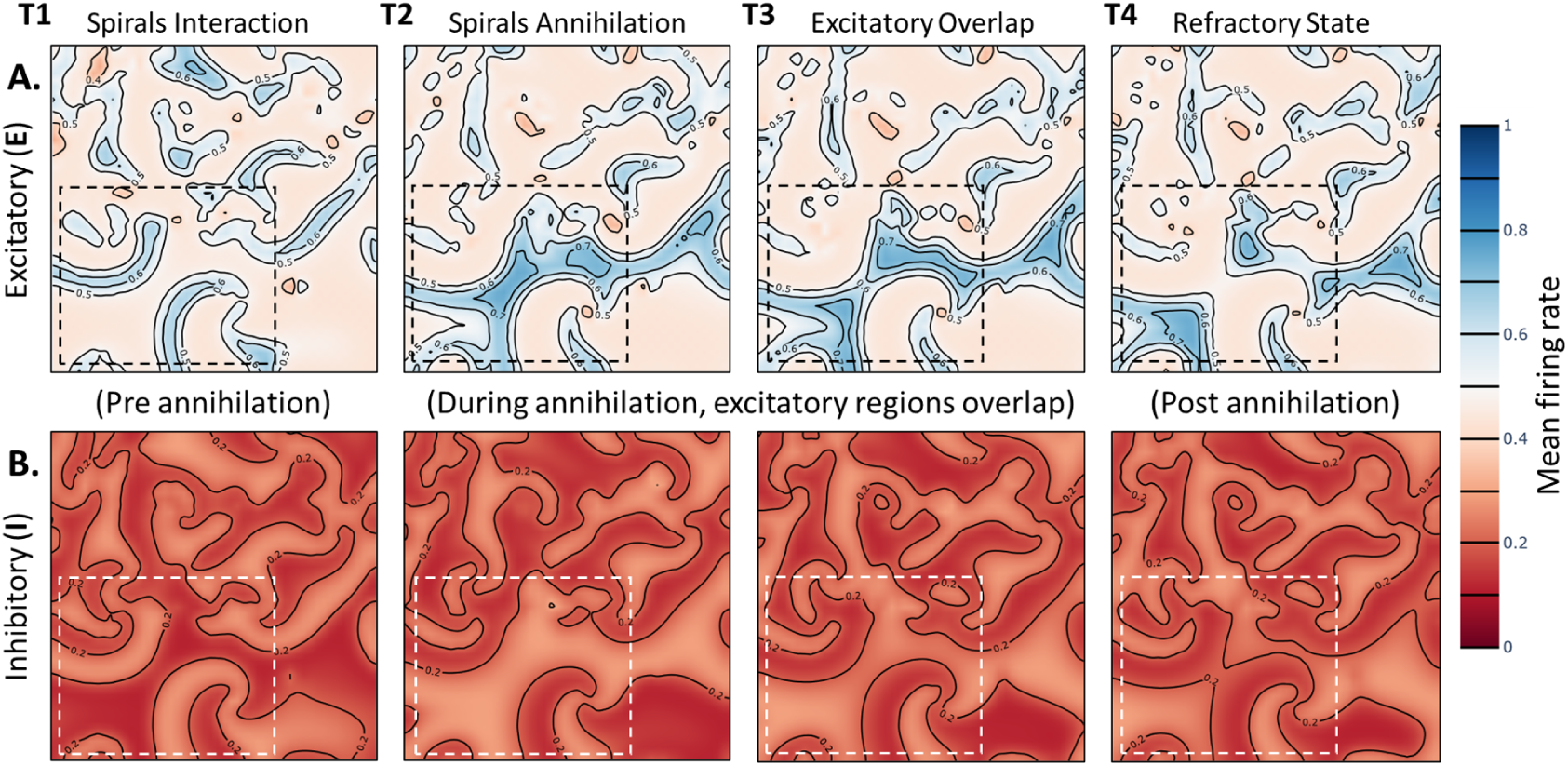
Annihilation of spiral traveling waves across the cortical fields. (A.) Spiral traveling waves propagating across the cortical excitatory (PYR) field with annihilation event shown within the black box. (B.) Annihilation events shown in the white box for the cortical inhibitory (PV) fields. Spiral arms originating from two different rotating spiral waves propagating towards each other at time (T1). These propagating patterns converge and annihilate each other at time (T2) resulting in the overlap of an excitation regions with slightly higher mean firing rate (T3). During post annihilation, these events diverge back into separate active regions (T4) entering into refractory state which prevents local excitation in the region for a small duration.

Occurrence of spontaneous traveling waves was shown using a 2D leaky integrate and fire (LIF) conductance based spiking network model considering distance dependent conduction delays (Davis et al., 2021). In current work, the rate model provides insight that the cortical circuits with local spiking activity as mean population dynamics with weak inhibitory coupling is sufficient enough for the emergence of complex spiral traveling waves. The prediction from the model that the complex spiral traveling waves are spontaneous in nature matches with previously proposed modeling studies (Davis et al., 2021). Here I also hypothesize that these distance dependent delays could also influence the inherent differences in the inhibitory time scales leading to formation of multiple slow and fast local systems across multiple cortical regions resulting in giving rise to mixed mode oscillatory behavior. A speculative idea is that these oscillations might give rise to different rhythms observed in the brain such as alpha, theta or beta, but the underlying mechanisms might be simple. An interesting aspect to investigate would be the concept of compression or reflection of cortical waves (W. Xu et al., 2007). The patterns of sharp waves ripples also undergo reflection (Patel et al., 2013). In the simulation results, if multiple local patterns were active together then how integration and compression of information could happens computationally for efficient working memory storage?

### Annihilation events across the Complex Spiral Traveling Waves

Annihilation events occur across the spatiotemporal cortical grid as seen in Figure 4. Two different spiral waves propagating towards each other at time (T1) annihilate each other at time (T2), resulting in the overlap of excitation regions at time (T3). After annihilation, spirals enter into a refractory state at (T4) which prevents local excitation in the region for a small duration. Spiral traveling waves propagates across the excitatory and inhibitory cortical field and undergoes annihilation. In addition, annihilation events, concentric activity along with spiral cores across the excitatory and inhibitory cortical field is described in more detail in the supplementary figure 2.

Complex spiral traveling waves in the cortex represent self-sustaining spatiotemporal patterns (Huang et al., 2004, 2010) as localized attractors that organize population dynamics across space and time. When two spirals annihilate, the system transitions from coexisting attractors (or nodes) to a unified state (source) or even to quiescence (sink). I hypothesize that a computational system might removes redundant or conflicting dynamic regimes to make a global “decision” when incompatible local waves collide and extinguish each other. It might reflect a reset mechanism of priors predictions in the predictive coding frameworks. In essence, annihilation may reflect controlled deletion of stored or encoded spatiotemporal information. Or might be a mechanism for winner-take-all selection which partitions the cortical tissue into coherent processing units. Annihilation could be a selective gating mechanism of stored information.

In excitable cortical media, the annihilation of spiral traveling waves represents a fundamental operation by which the system resolves competing spatiotemporal patterns. Each spiral carries phase-encoded information and defines a localized dynamical attractor, their mutual annihilation constitutes both the erasure and the updating of cortical information states. This event effectively enforces winner-take-all selection among incompatible population patterns, thereby enabling segmentation, decision-making, and resetting of local computations. Moreover, annihilation plays a stabilizing role by preventing runaway propagation and maintaining the cortex in a flexible regime near criticality. Thus, spiral wave annihilation is not simply a dynamical artifact, but an essential mechanism for regulating information flow, preserving stability, and enabling reconfiguration of cortical computation (Chemla et al., 2019).

### Weak Grating Stimulus across the Cortical Field

Now, after investigating and hypothesizing the possible role of annihilation events as selective information gating mechanisms of high frequency local messenger waves for slow frequencies traveling carrier waves, it seems intriguing to look at how the cortical grid responds to grating stimulus. A 2D network of bidirectionally coupled pyramidal neurons representing a cortical sheet in the hippocampus, where an adaptive EIF model described a single neuron, could produce spiral wave patterns (Souza et al., 2024). A similar finding proposed about the effect of weak or a strong external stimulus depending on the amplitude and duration modulates the underlying spiral patterns might results in plane waves propagation in the network (Li et al., 2025). Similar finding is observed in the simulation results for the cortical excitatory field shown in the figures 5, and 6. It also matches with the preliminary raw patterns for more details in the supplementary figure 3. Interestingly, the single layer based on a chaotic neural network (Aihara et al., 1990), which is composed of mutually coupled Aihara chaotic neurons, and the spiral patterns dynamics appear very similar to those described in the supplementary figure 2. These predictions clearly show that a grating stimulus’s properties could determine the working memory characteristics for encoding and maintenance of information.

**Figure 5:**
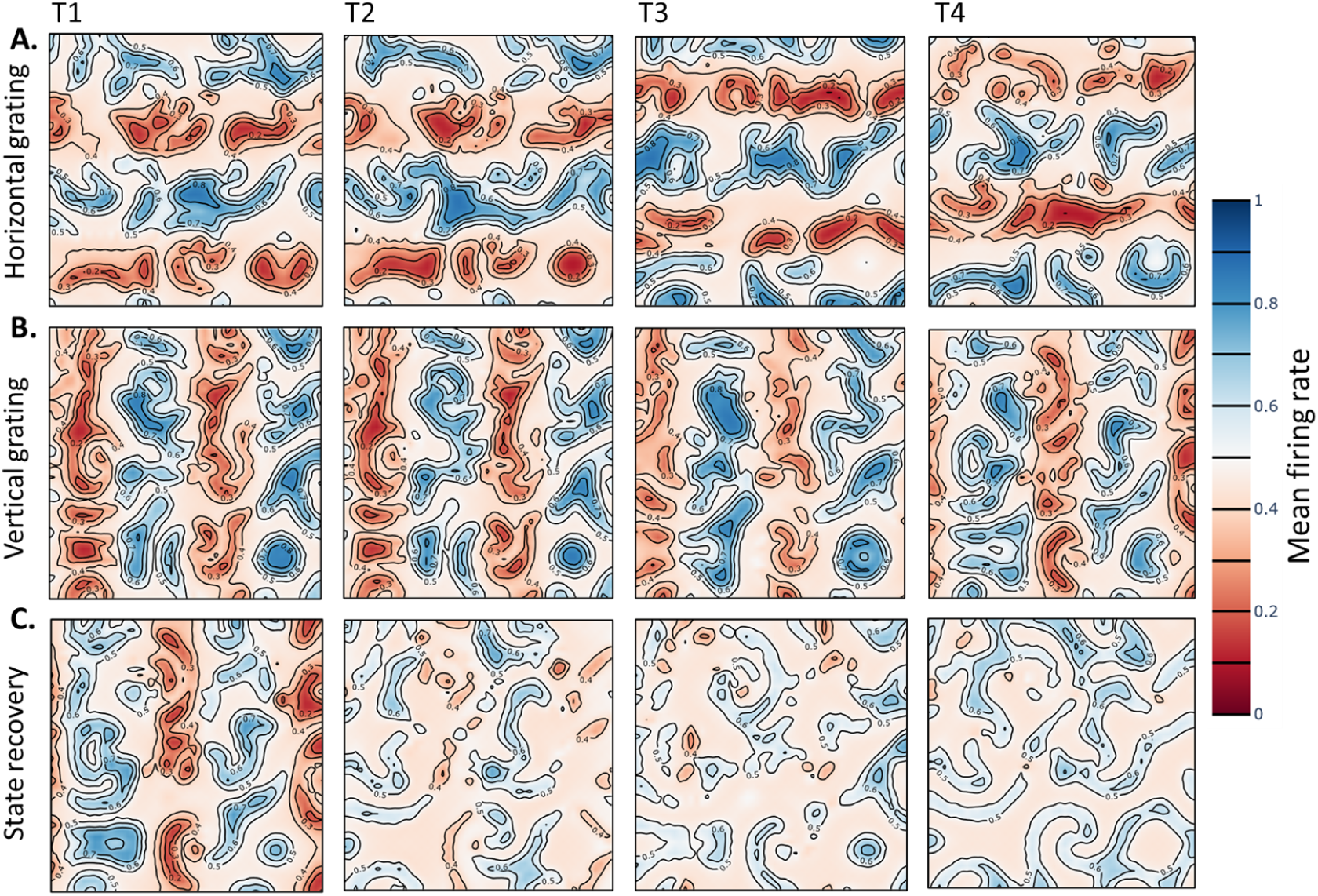
Effect of a weak grating stimulus across the cortical spiral field. (A.) A weak grating stimulus with small amplitude is applied in the horizontal direction across an excitatory cortical field. (B.) Similarly, a weak grating stimulus with small amplitude is applied in the vertical direction across an excitatory cortical field. During the time from (T2-T4), these externally applied grating patterns exists for a small duration of time as transiently modified spirals patterns exists across the state space. (C.) Over the time, these transient patterns (or states) recover back to the original spiral patterns before the application of external stimulus displaying short term memory effect behaviors.

As seen in the figure 5, a weak stimulus with small amplitude is applied in the horizontal direction across an excitatory (PYR) spiral cortical field. In the fig 5B., similarly, a weak grating stimulus with small amplitude is applied in the vertical direction across an excitatory spiral cortical field for a small duration. During the time interval between (T2-T4), these externally applied grating patterns exist for a short duration even after the removal of the grating stimulus, as transiently modified spiral patterns spanning the state space in fig. 5C. Over slightly longer duration, these transient patterns (or states) recover back to the original spiral patterns seen before the application of an external stimulus, displaying memory effect. Hence, its a compelling evidence to describe this effect as a *cortical working memory*.

### Strong Grating Stimulus across the Cortical Field

Previously applied weak grating stimulus to the cortical field resulted in the recovery of original states from transiently modified states. Forgot to mention that the idea of applying external stimulus was inspired from Huber and Wiesel classical experiments that utilized grating stimulus inputs to test response of receptive field (Hubel & Wiesel, 1959). In the figure 6, a strong grating stimulus with large amplitude is applied at 120 degree (appearing as negative slope) in the slanted direction in fig. (6A.) or in the vertical direction in fig. (6B.) across an excitatory (PYR) cortical field. Further, stimulus application results in the existence of memory patterns for a short time post-stimulus, which might undergo encoding or maintenance of the visual stimulus via modified spiral patterns, suggesting working memory characteristics. Patterns or traces may gets encoded or maintained across the inhibitory (PV) cortical field or its traces seen in the fig. (6C.). These complex spiral waves patterns undergo state modification which could be seen by comparing cortical states before and after the application of a grating stimulus across an excitatory cortical field in fig. (6D.). Similar comparison reflects cortical states before and after the application of a grating stimulus across an inhibitory cortical field in fig. (6E.). Its clearly visible on comparison that the mean firing rate of an excitatory cortical field is greater than that of an inhibitory field. But the regions are more discrete and discontinuous in PYR regions compared to smooth and low amplitude inhibitory cortical field regions.

**Figure 6:**
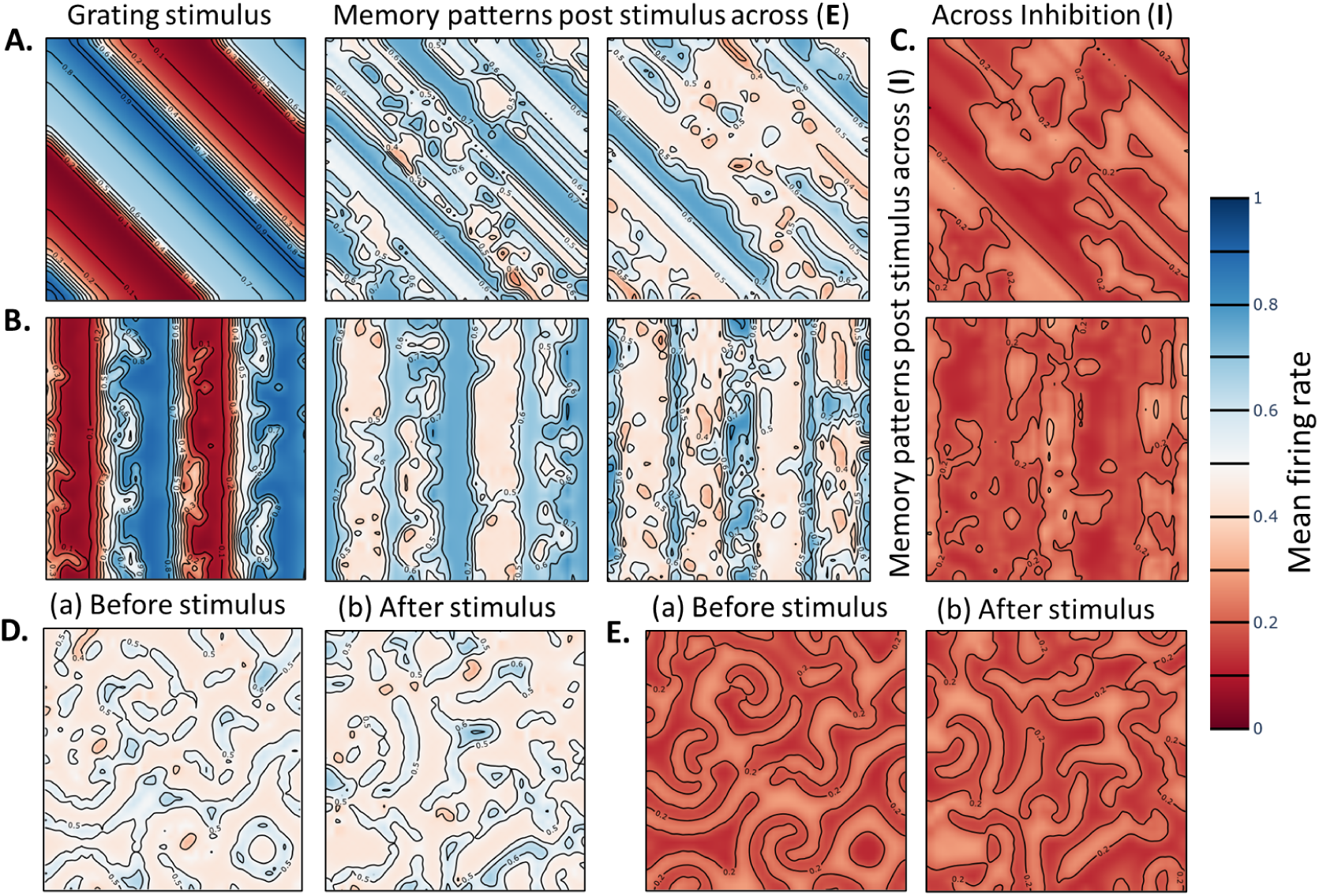
Effect of a strong external grating stimulus across the cortical spiral field. A strong grating stimulus with large amplitude is applied in the slanted direction (A.) or in the vertical direction (B.) across an excitatory (PYR) cortical field. Further it results in the existence of memory patterns for a short duration of time post stimulus which might undergo encoding or maintenance of such visual stimulus via modified spiral patterns suggesting working memory characteristics. (C.) Representation of the patterns encoded or maintained across the inhibitory cortical field. Complex spiral waves undergoes state modification. (D.) Comparison of cortical states before and after the application of grating stimulus across an excitatory cortical field. (E.) Comparison of cortical states before and after the application of grating stimulus across an inhibitory cortical fields.

### Phase Gradient and Velocity Field for Complex Spirals Traveling Waves

The phase *ϕ*(*x, t*) was extracted via the Hilbert transform of the spatially distributed signal, capturing the local timing of population oscillations. Complex spiral waves exhibited pronounced topological heterogeneity with multiple singularities—points where the phase is undefined and rotates by 2*π*-identified using circular integrals around minimal closed loops to distinguish clockwise and counter-clockwise cores. Local phase-gradient vectors quantified the instantaneous direction and velocity of wave propagation (Townsend & Gong, 2018), with steep gradients marking rapid propagation or collision fronts and shallow gradients indicating slower or disorganized flow. This spatiotemporal analysis captures metastable transitions as singularities drift, annihilate, or re-emerge, showing that complex spiral waves evolve through transient, partially ordered configurations shaped by recurrent connectivity, mesoscale heterogeneity, and intrinsic neural fluctuations, linking large-scale traveling waves to flexible cortical coordination. Phase gradient fields are shown in figure 7. The phase dynamics of the excitatory cortical population are rapid, revealing the underlying spontaneous neuronal activity patterns with great temporal detail. In contrast, the inhibitory cortical population exhibits smoother propagation of the rotating spirals over time. Phase gradients along with phase-velocity cortical fields for an excitatory and an inhibitory population are shown in the supplementary video (3,4,5) file.

**Figure 7:**
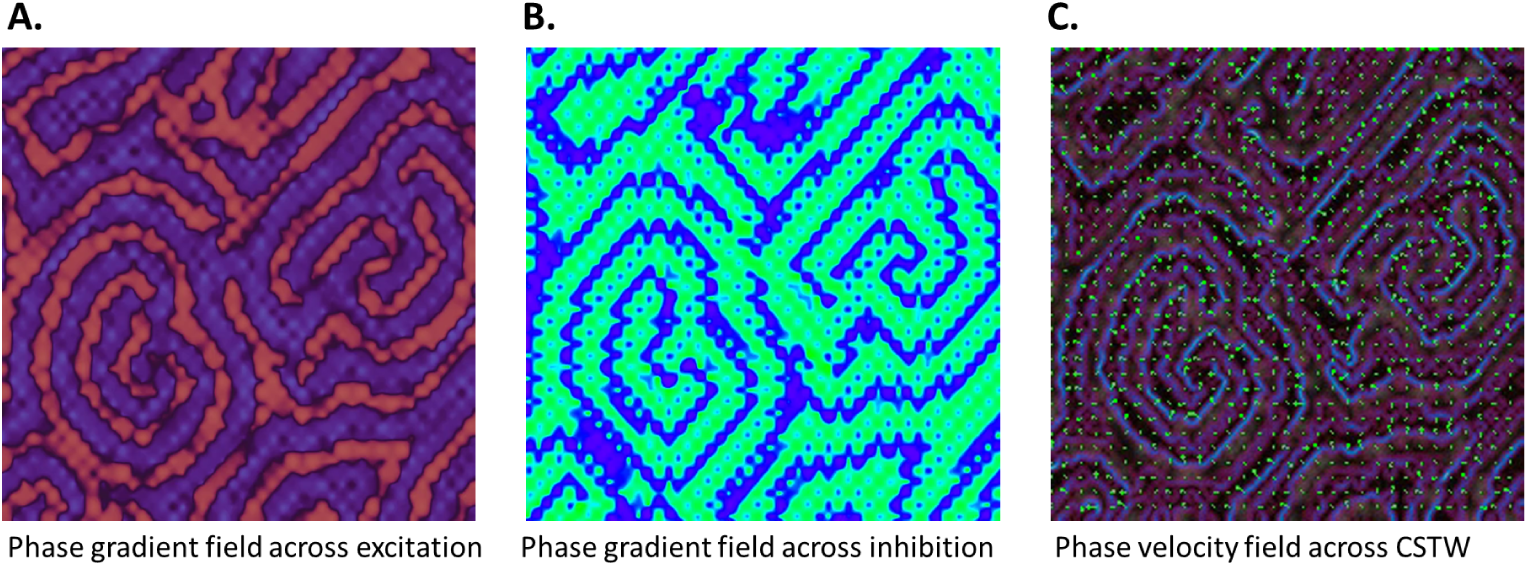
Phase gradient and velocity field across complex spiral traveling waves. (A.) Phase gradient field across the PYR. (B.) Phase gradient field across the PV. (C.) Phase velocity field with quiver plot.

The phase vector field represents the local direction of the traveling wave at each point in space. Instantaneous phase at each spatial location is determined using the Hilbert transform and then computing the gradient of the phase. Useful to identify the core of spiral waves also knows as phase singularities points where the phase is undefined. Further using topological analysis, different traveling wave patterns such as spiral, planar, concentric or complex multi spirals were detected. The phase velocity field tells you the speed and direction of the local phase propagation. Essentially, it’s the temporal change of the phase at each point. Essentially, it shows how fast the waves move across the cortical sheet. It helps to quantify the wave speed and measures how fast spiral arms or wavefronts travel. Second, detects dynamic changes as spiral waves can accelerate, decelerate, or meander; phase velocity field captures this in real time. Third, most important for complex spiral waves is that the variations in the phase velocity can indicate heterogeneity in the cortical excitability which we observed shown in the supplementary video. Both the gradient and velocity fields provide complete characterization of spiral waves as phase vector field tells where the wave is going (direction) and the phase velocity field tells how fast it is moving (speed).

## Material and Methods

### Cortical Field Model

A spatially extended Wilson-Cowan based neural field model is proposed referred as *cortical field model ^1^ [^1^No experimental data were used for the model, all results are simulations based.]* as it contains four different interacting cortical neural populations defined over a two-dimensional grid with periodic boundary conditions. The populations consist of an excitatory pyramidal (PYR) population as *E*(*x, y, t*) and three distinct inhibitory types namely fast-spiking parvalbumin (PV) as *I*_1_(*x, y, t*), dendrite-targeting somatostatin (SOM) as *I*_2_(*x, y, t*), and disinhibitory (VIP) as *I*_3_(*x, y, t*). Model evolves according to a reaction-diffusion with non linear coupling using euler integration. Each neural population (N) dynamics evolving over the 2D spatiotemporal field over time follows:

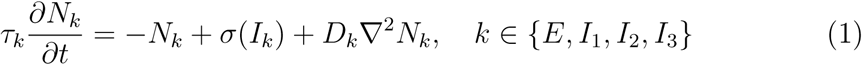

where *N* (*x, y, t*) denotes the local activity field, *σ*(·) is the nonlinear sigmoidal activation function that transforms the synaptic input into firing rate, *I_N_* (*x, y, t*) = *WN* + *P* is the net synaptic input to population *k*, *W* is the effective recurrent connectivity matrix, *P* (*x, y, t*) represents external spatio temporal input, *τ* is the population-specific intrinsic mean firing rate constant, and *D* is the diffusive coupling coefficient.

### Extended Cortical Grid Model

Why reaction diffusion formalism for the cortical tissue? Local non linear interactions through excitatory and multiple inhibitory dynamics can be captured simplistically using wilson cowan formalism that defines the local interactions. Whereas long range connections among the populations with distance dependent delays approximated by diffusion (isotropic or anisotropic spread of activation). Brain activity combines nonlinear local excitation/inhibition with distant long range spatial spread of activity through axonal delays which naturally gives rise to wave phenomena (traveling fronts, oscillations, spirals), and these patterns are experimentally observed across cortex and retina. Next, mixed-mode oscillations (MMOs) and spiral rotating waves (SRWs) are both complex dynamical behaviors that arise in nonlinear excitable or oscillatory systems, particularly in reaction-diffusion systems, electrochemical systems, neuronal models, and chemical oscillators like the Belousov-Zhabotinsky reaction. MMOs are temporal oscillations characterized by alternating small and large amplitude oscillations (SAOs and LAOs). Typically arise in slow-fast systems due to canard dynamics, folded node singularities, or delayed Hopf bifurcations. SRWs are spatial patterns that rotate around the core and occur in reaction-diffusion systems with excitable or oscillatory kinetics. While they may appear different, MMOs being temporal phenomena and spirals being spatiotemporal, they can be intimately related through the underlying dynamical systems framework, especially in slow-fast systems with multiple timescales. Reaction-diffusion systems can exhibit slow-fast behavior in both time and space. The core of the spiral wave and its surrounding arms can correspond to regions where MMOs emerge in the local temporal dynamics. The tip of a spiral often exhibits complex dynamics including quasi-periodicity and even MMOs. The periodic alternation of fast activations and slow recovery phases at the tip can mirror an MMO structure in time. Hence, an extened version of the rate model to capture these phenomena. The synaptic input to each population is modeled as a weighted sum of activities from all other populations, combined with an external input:

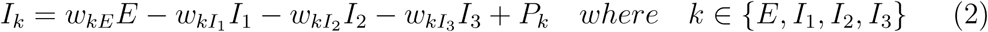

where *w_ij_* denotes the synaptic weight from population *j* to *i*, and *P_i_* is a constant external input. The transformation from synaptic input to firing rate is governed by a sigmoidal function:

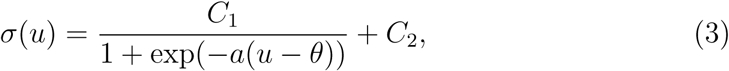

where *a* is the gain, *θ* is the threshold, and *C*_1_*, C*_2_ control the range and baseline firing rate output.

The spatial coupling or spread of the activity via diffusion is modeled via the Laplacian operator, discretized using a 5-point stencil with periodic boundary conditions:

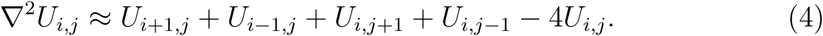

This implements isotropic diffusion across a regular 2D grid with homogeneous local connectivity. The numerical integration system is integrated in time using an explicit forward Euler scheme:

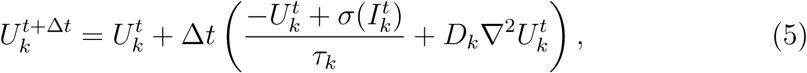

where Δ*t* is the simulation time step, and *k* indexes each population. All updates are performed in-place for computational efficiency, and simulations are typically run on a grid of size *N* × *N* with periodic boundary conditions in both spatial dimensions.

The external grating stimulus is defined as

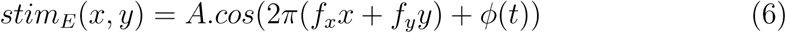

where A is the amplitude, (*f_x_, f_y_*) are spatial frequencies in x and y directions, *ϕ*(*t*) is the phase.

The model is governed by 28 synaptic and external input parameters, and four diffusion coefficients. Time constants: *τ_E_, τ_I_*_1_ *, τ_I_*_2_ *, τ_I_*_3_ , Synaptic weights: *w_kl_* for *k, l* ∈ {*E, I*_1_*, I*_2_*, I*_3_}, external inputs: *P_E_, P_I_*_1_ *, P_I_*_2_ *, P_I_*_3_ , sigmoid parameters: *a, θ, C*_1_*, C*_2_, diffusion coefficients: *D_E_, D_I_*_1_ *, D_I_*_2_ *, D_I_*_3_ . Parameter values are selected to preserve stability and to match biologically plausible firing regimes. Parameter sweeps are conducted to explore the impact of coupling strengths and external drive on emergent spatiotemporal patterns.

All simulations were performed on a two-dimensional *N* × *N* grid with periodic boundary conditions. Unless otherwise stated, *N* = 100 and the spatial grid spacing *h* = 1 was used. The time step for numerical integration was set to Δ*t* = 0.1 ms, ensuring numerical stability for the explicit Euler scheme. Initial conditions were given by a small-amplitude spatially uncorrelated Gaussian noise to each population:

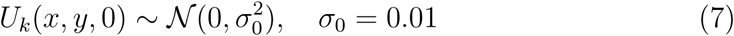

Each simulation ran for a longer duration of 3000 ms. After discarding an initial 1000 ms transient, initially random patterns occurs and eventually over time spiral patterns emerges, I analyzed these pattern formation, spatial synchrony, and temporal oscillations. Local cortical circuit rate-model dynamics were simulated in Python, and XPPAUT (B. Ermentrout & Mahajan, 2003) was used to compute bifurcation diagrams. All numerical experiments were implemented in NumPy with vectorized operations for efficiency, and visualizations were generated using Matplotlib. Numba and a Dash application were employed to enable interactive, real-time visualization along with parameter exploration.

### Relation to Prior Models

Classical neural field models typically assume a single, homogeneous inhibitory population and therefore miss important interactions that drive emergent cortical dynamics. Here we extend the Wilson–Cowan framework (Wilson & Cowan, 1972) and its reaction–diffusion variants (Amari, 1977; Bressloff et al., 2001; B. Ermentrout, 1998) by introducing three inhibitory subtypes with distinct synaptic and dynamical properties. This added diversity, motivated by physiological evidence of interneuron heterogeneity (Kepecs & Fishell, 2014; Tremblay et al., 2016) and recent multi-population models of working memory (Murray et al., 2014) enables the spatially continuous system to generate structured spatiotemporal patterns that cannot arise in traditional two-population formulations.

## Discussion

Here, a four-population cortical field model that extends classical excitatory-inhibitory frameworks by incorporating three distinct inhibitory subtypes: fast-spiking (PV-like), dendrite-targeting (SST-like), and disinhibitory (VIP-like) interneurons is presented. Through simulations, it was shown that this local cortical circuit with multiple time scale dynamics could produce mixed mode oscillations with small and large amplitude oscillations representing existence of multiple rhythms. Further extending this local circuit coupled by diffusion representing the axonal delays forming a spatiotemporal 2D cortical grid shows emergence of complex rotating spiral traveling waves and other patterns such as concentric, planar, source or sink seen experimentally.

Based on the findings, I hypothesize that the annihilation of two or more cortical spirals could corresponds to erasure of stored spatiotemporal information, computationally. There is no direct empirical or modeling study that shows cortical spiral wave annihilation as *“information deletion”*. Here, I speculate that the cortex uses wave annihilation as a *“decision”* mechanism between competing dynamical states. Annihilation events might implement a resets of local modules or clearing prior state as in predictive coding, again this is a hypothesis, not backed by data. Again, I make a reasonable proposal that spiral wave annihilation may be a dynamical integration or an erasure mechanism in cortical circuits. When incompatible spirals waves collide, their mutual annihilation removes the encoded representations from the local substrate, producing a transient reset. While annihilation itself destroys the prior pattern (a deletion operation), it creates the conditions for subsequent integration by freeing cortical resources during working memory, opening a temporal window for new inputs to impose coherent activity. Thus, annihilation could implements selective gating mechanisms. This hypothesis is based on the observations that complex spiral waves rotating in clockwise or anticlockwise direction in the cortex, carry spatiotemporal phase information on collisions do lead to annihilation. Hence, it’s reasonable to propose that on waves annihilation something like an *“information erasure”* event might occur.

Traditional two-population models consisting of a single excitatory and a single inhibitory population offer insights into fundamental mechanisms such as excitation inhibition balance and cortical pattern formation. However, recent anatomical and physiological studies have revealed that cortical inhibitory neurons comprise multiple distinct subtypes with specific connectivity profiles, intrinsic time constants, and modulatory roles. In particular, parvalbumin-positive (PV), somatostatin-positive (SST), and vasoactive intestinal peptide-expressing (VIP) interneurons exhibit markedly different synaptic targets and temporal dynamics. While our model captures essential features of interneurons diversity, several limitations remain. First, firing-rate equations, which abstract away spiking dynamics, synaptic delays, and conductance nonlinearities. Incorporating spiking network models or conductance-based dynamics would yield more biologically realistic insights. Second, plasticity or learning is not modelled, future work could explore how inhibitory subtype-specific plasticity shapes pattern formation. Third, different topologies of the local cortical circuits are not tested which might display unique regimes of operations such as global synchronization, oscillations or different patterns. Additionally, stability analysis or rigorous bifurcation theory on the mechanistic local cortical circuits can complement the results and provide deeper insight into regime transitions. Here a 2D spatiotemporal cortical grid model is proposed. But the cortex in a 3D structure. It would be interesting to investigate how does the brain utilizes these different types of cells across the cortical layers with distinct connectivity by developing a 3D cortical field model.

In summary, the cortical complex spiral traveling waves emerge due to inhibitory weak coupling between local cortical circuits composed of an excitatory (PYR) and distinct inhibitory neural populations (PV, SOM, and VIP) across a 2D cortical grid. Key findings suggest that these complex spiral waves are regulated by local interactions, such as sources and sinks, that appear planar or concentric within local circuits. Mixed-mode oscillations (MMOs) may underlie the robust coexistence of multiple rhythms despite synaptic heterogeneity across cortical layers. Overall, the results emphasize the critical role of interneurons diversity in shaping cortical dynamics. By integrating multiple inhibitory populations into spatially extended rate models, I uncover novel mechanisms by which local interactions give rise to global brain activity patterns. This framework opens new directions for linking microcircuit physiology to macroscopic brain dynamics observed experimentally to link mesoscopic activity with macroscopic dynamics. Further, the cortical grid model provides a possibility to develop more complex scalable layered cortical architecture in the near future.

## Acknowledgment

This research was conducted independently outside the scope of the author’s institutional duties and received no external funding. No experimental data were used for the work. The author conceptualized, developed the computational framework, performed simulations and analysis, interpreted the results, and wrote the manuscript. The views expressed in this work are solely those of the author.

## Supplementary Figures

**Supplementary Figure 1:**
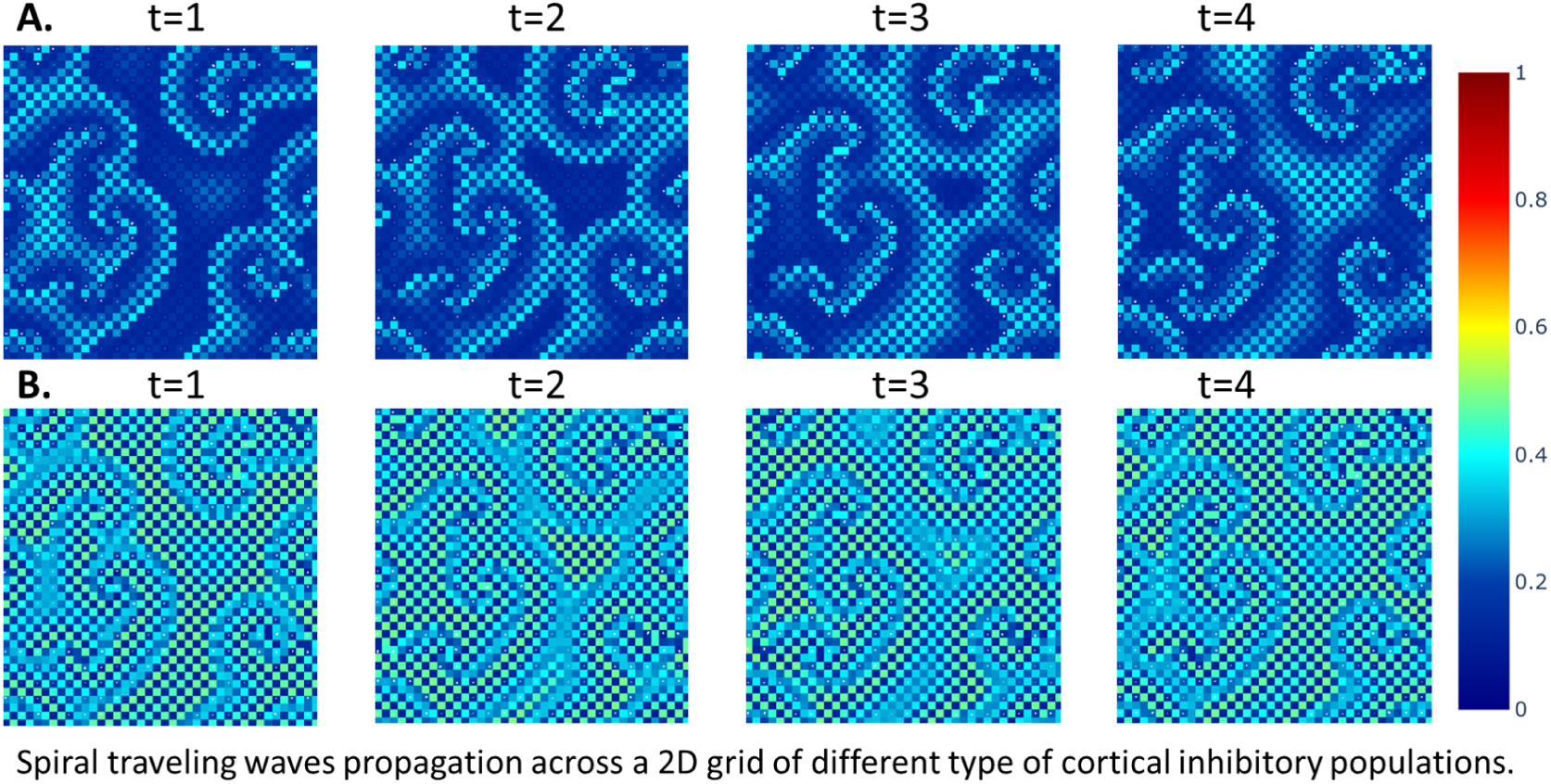
Spiral traveling wave propagation across cortical inhibitory field. (A.) Propagation of spiral patterns across SOM inhibitory field with low amplitude at different time steps t=(1-4). In (B.), spiral propagation across PV inhibitory field t=(1-4). Propagation is smoother across cortical inhibitory fields.

**Supplementary Figure 2:**
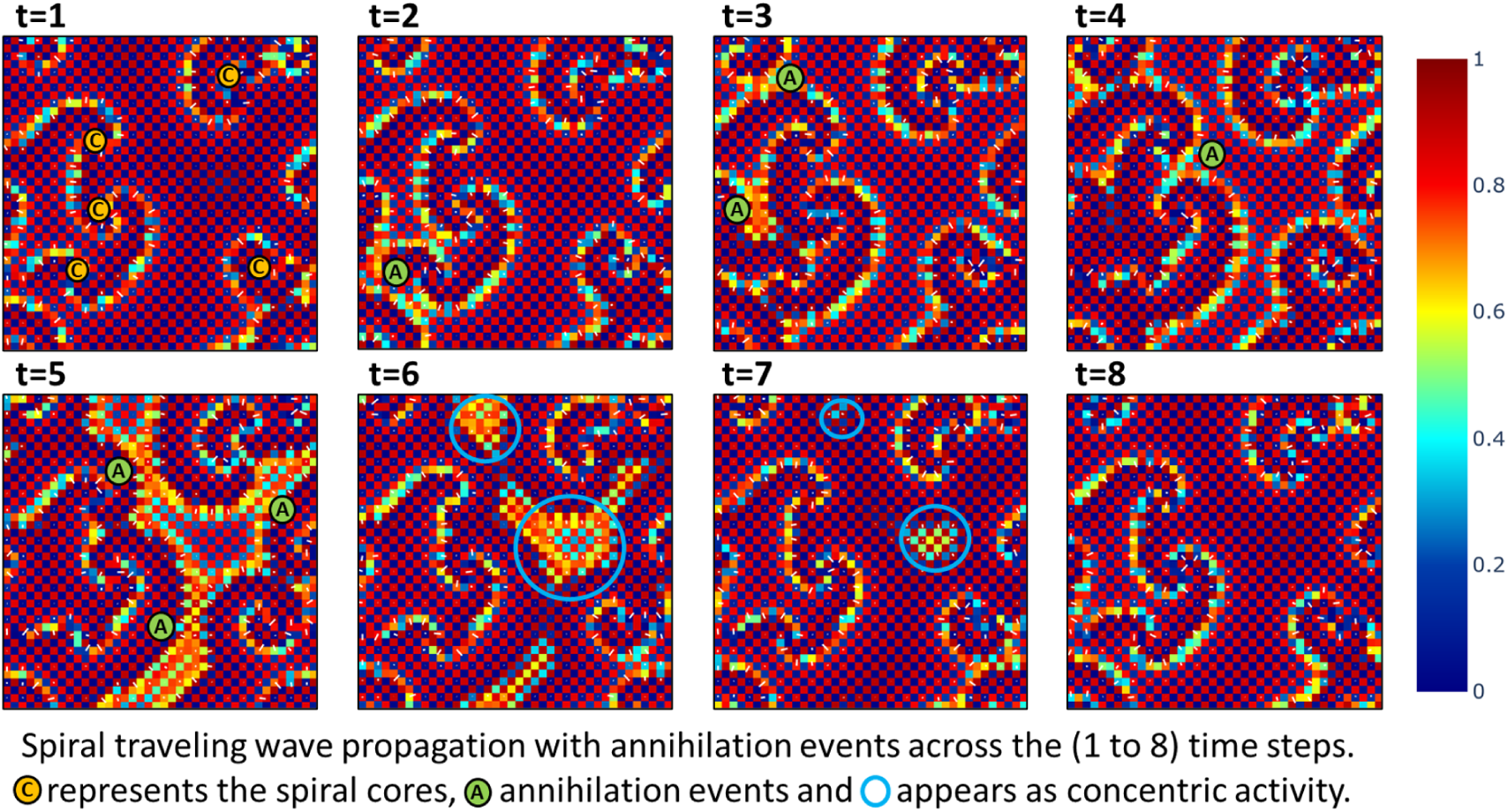
Spiral traveling wave propagation across the cortical fields. Spiral patterns propagate across cortical excitatory (PYR) fields. Annihilation events propagate across the time steps t=(1-8). Annihilation events are shown with filled green circle (A), spiral cores in orange filled circle (C), and somewhat concentric or convergence of annihilation is shown in empty blue circle. Activity is quite complex and heterogeneous in nature.

**Supplementary Figure 3:**
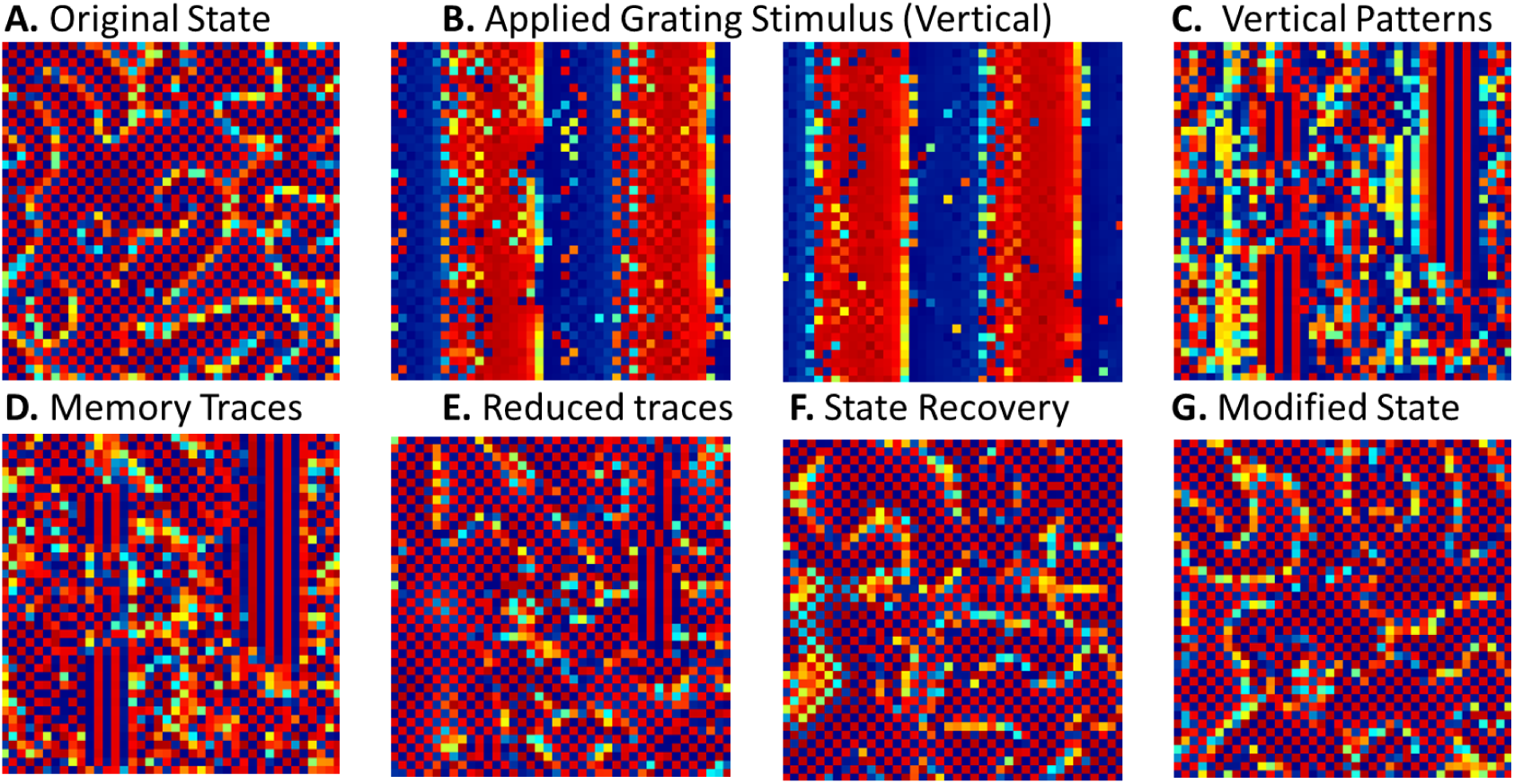
Effect of applying a grating stimulus on the spiral traveling waves. Spiral traveling wave modulation based on an applied vertical grating stimulus (B) results in vertical patterns (C) persists over time post stimulus showing short term memory effect (D). These memory traces vanishes (E) during the state recovery (F) back to a modified state (G)

**Supplementary Figure 4:**
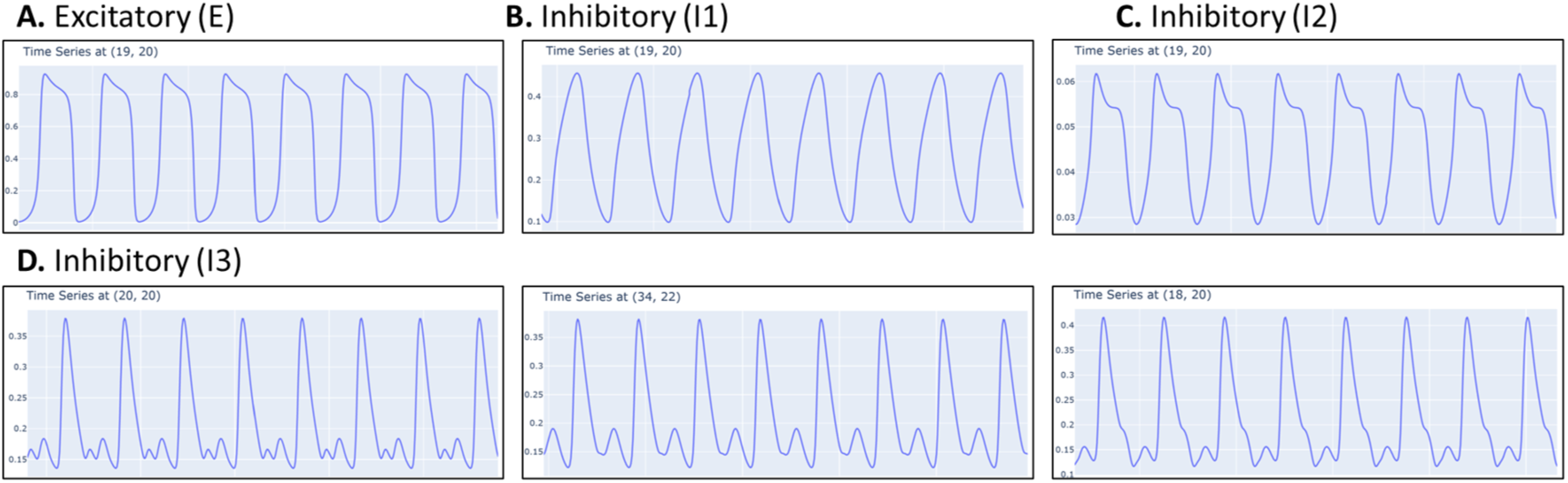
Complex spiral traveling waves time series data. Time series data across different grid points for excitatory and different inhibitory types (*I*_1_*, I*_2_*, I*_3_). In (A.,B., and C.), time series across the grid point shows normal oscillations. In (D.), time series shows mixed mode oscillations (MMOs) with large amplitudes oscillations (LAOs), two small amplitude oscillations (SAOs) in the first case and one SAOs in second and third case.

**Supplementary Figure 5:**
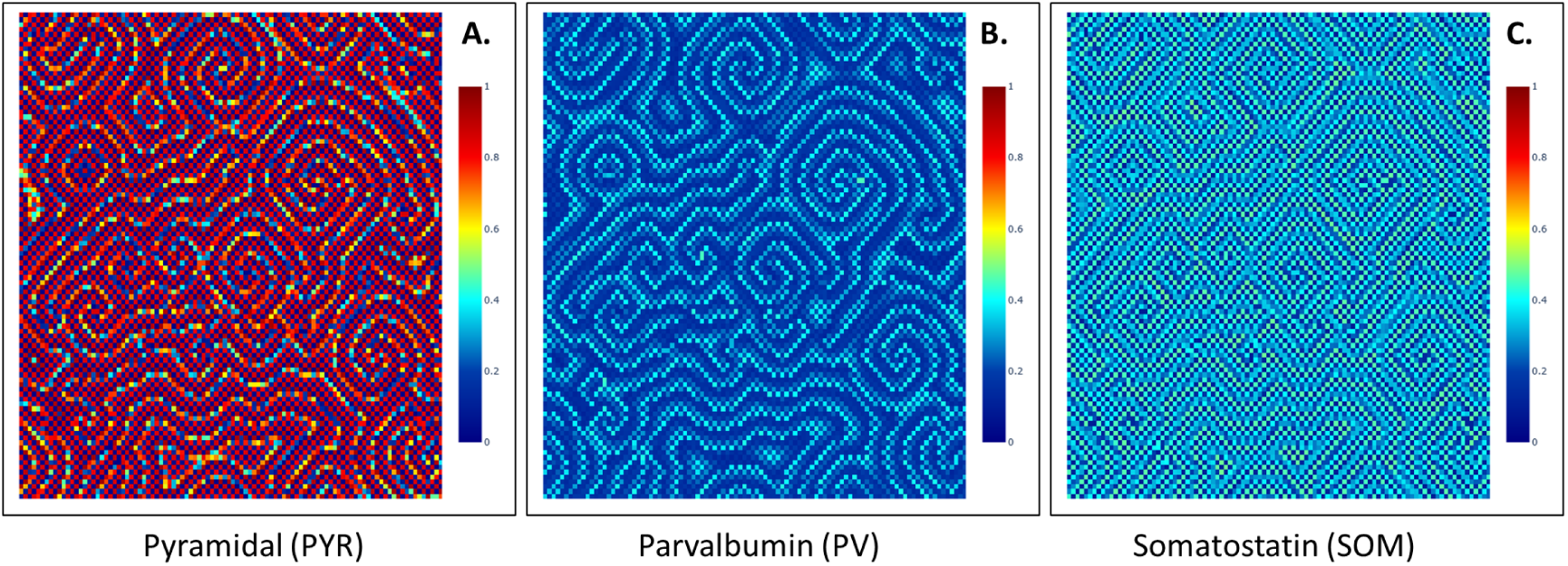
Multiple spiral traveling waves across the 2D cortical grid (100x100) (A.) Many complex spirals rotating in both clockwise (CW) and anticlockwise (ACW) directions in the PYR field (B.) Spirals with small amplitude and propagation speed occurs across PV and (C.) SOM.

**Supplementary Figure 6:**
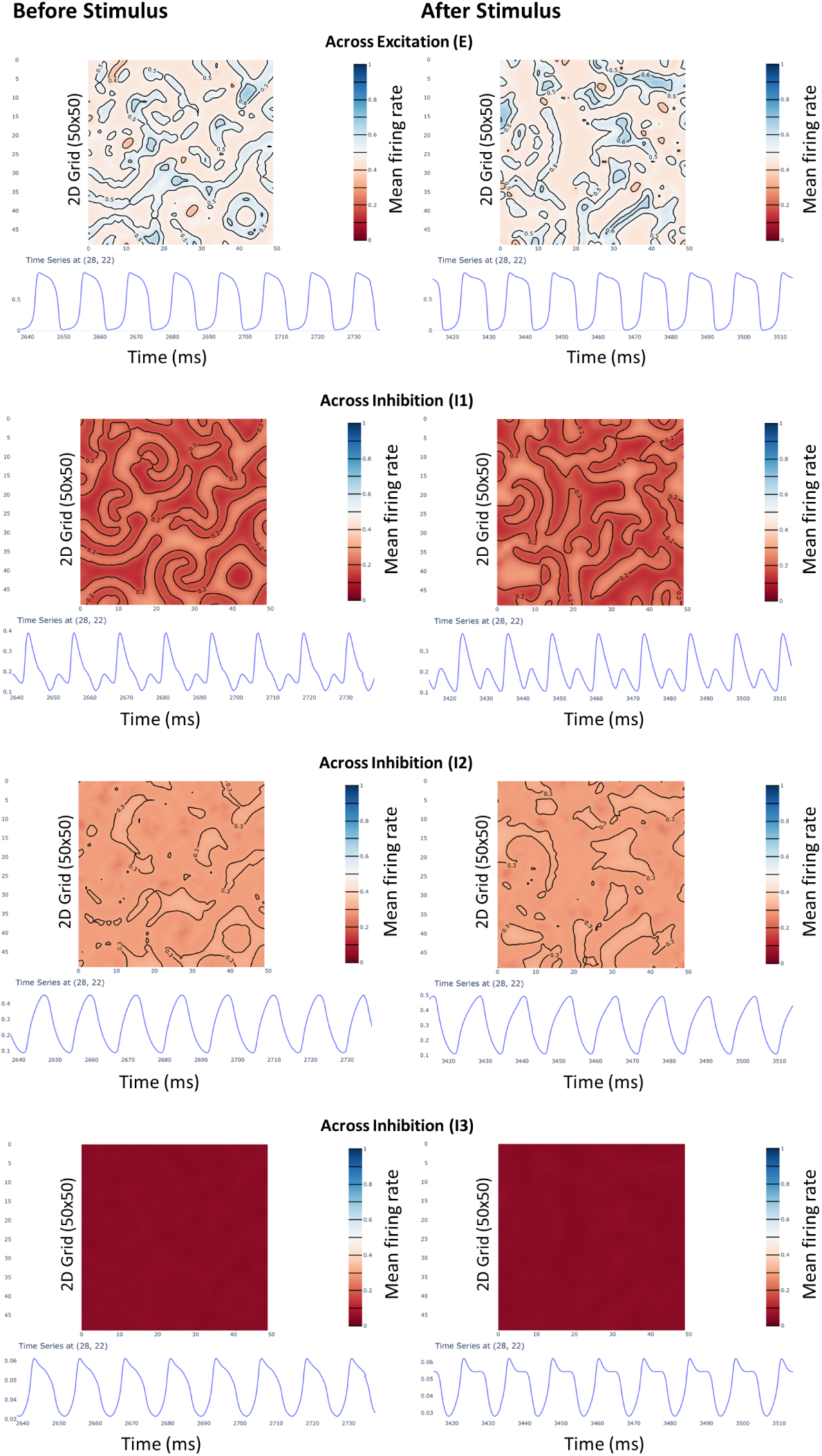
Effect of grating stimulus on the Spiral States. Spiral cortical field state before and after the grating stimulus across PYR, PV, SOM and VIP. Also, as seen in the time series at a particular grid point (28,22) is modified. Multiple rhythms coexists together.

